# Characterizing the role of Ca^2+^ fluxes in defining fast and slow components of spontaneous Ca^2+^ local transients in OPCs using computational modeling

**DOI:** 10.1101/2022.06.02.494570

**Authors:** Nicolas Desjardins, Xiaoyu Jiang, Laurentiu Oprea, Kushagra Sareen, James Q. Zheng, Anmar Khadra

## Abstract

Spontaneous Ca^2+^ local transients (SCaLTs) in isolated oligodendrocyte precursor cells (OPCs) are largely regulated by the following fluxes: store-operated Ca^2+^ entry (SOCE), Na^+^/Ca^2+^ exchange (NCX), Ca^2+^ pumping through Ca^2+^-ATPases, and Ca^2+^-induced Ca^2+^-release through Ryanodine receptors (RyR) and inositoltriphosphate receptors (IP_3_R). However, the relative contributions of these fluxes in mediating fast spiking and slow baseline oscillations seen in SCaLTs remain incompletely understood. Here, we developed a stochastic spatiotemporal computational model to simulate SCaLTs in a homogeneous medium with ion flow between the extracellular, cytoplasmic and endoplasmic-reticulum compartments. By simulating the model and plotting both the histograms of SCaLTs obtained experimentally and from the model as well as the standard deviation of interspike intervals (ISI) against ISI averages of multiple model and experimental realizations we revealed that: SCaLTs exhibit very similar characteristics between the two datasets, they are mostly random, they encode information in their frequency, and the slow baseline oscillations could be due to the stochastic slow clustering of IP_3_R (modeled as an Ornstein-Uhlenbeck noise process). Bifurcation analysis of a deterministic temporal version of the model shows that the contribution of fluxes to SCaLTs depends on the parameter regime and that the combination of excitability, stochasticity, and mixed-mode oscillations are responsible for irregular spiking and doublets in SCaLTs. Additionally, our results demonstrate that blocking each flux reduces SCaLTs frequency and that the reverse (forward) mode of NCX decreases (increases) SCaLTs. Taken together, these results provide a quantitative framework for SCaLT formation in OPCs.

## 2 Introduction

The oligodendrocyte lineage of cells is responsible for myelinating axonal fibers in the central nervous system (CNS) [25]. Oligodendrocyte progenitor cells (OPCs, also known as NG2 cells) develop into immature oligodendrocytes (imOLs) and finally into myelinating mature oligodendrocytes (OLs), with each developmental class having distinct roles in the creation of white matter [5, 33]. Evidence suggests that some control of myelination must depend on neuronal activity [17, 9, 13, 12]. This means that during development, strict energy budgets dictate how much myelin is allocated to each neuron [35].

According to the adaptive myelination hypothesis, it has been suggested that if neural electrical activity regulates subcellular events in OPCs necessary for myelin elaboration, then myelin would form preferentially on the electrically active axons [4, 23]. Evidence in favour of this hypothesis was first found in the electrical activity-induced vesicular release by neurons, leading to increases in the number of rat optic nerve OPCs [2]; it was later experimentally induced by stimulating cortical layer V projection neurons, using optogenetics, causing OPCs in the deep layers of the premotor cortex and projections through the corpus callosum to proliferate [13]. Interestingly, within the stimulated premotor areas both the number of OLs and myelin thickness were increased.

Ca^2+^ signalling is the main pathway by which neural electrical activity mediates changes in myelination in OPCs. It is the key signalling molecule implicated in activity-dependent myelin growth in OPCs [27, 1, 24] and in controlling Stim1 and Golli, the two proteins in the myelin basic protein (MBP) family interactome [30]. Inhibition of Ca^2+^ transients in OPCs impairs not only their process elaboration and branching, but also myelin sheath size [32] and growth cone control [34]. Indeed, slow, high-amplitude somatic Ca^2+^ currents cause greater diffusion of somatic Ca^2+^ to myelinating processes [37].

The frequency of Ca^2+^ transients appears more important than the absolute amplitude of fluctuations. Rate modulating experiments showed that Ca^2+^ induced myelin sheath elongation (in imOLs) is promoted by high frequency Ca^2+^ transients [18]. On the other hand, low frequency Ca^2+^ transients (or long Ca^2+^ bursts) promote shortening [18]. However the average rate of transients depends on cell maturity. Immature oligodendrocytes have a high frequency of Ca^2+^ transients, and the frequency of such transients increases as OL processes grow and elaborate [3].

Interestingly, spontaneous Ca^2+^ local transients (SCaLTs) are also present in isolated OPCs (i.e., in the absence of synaptic inputs from neurons), with profiles that exhibit both fast spiking along with underlying slow oscillations [32]. This highlights the ability of OPCs in spontaneously generating such activity intrinsically. Evidence suggests that SCaLTs are produced by Ca^2+^ mobilization due to (i) Ca^2+^ entry into the cytosol governed by three main fluxes including store-operated Ca^2+^ entry (SOCE) using three primary SOCE channel proteins: ORAI1, STIM1, and STIM2 [30, 32], Na^+^/Ca^2+^ exchangers (NCX) [3, 6] that exchange three Na^+^ ions for one Ca^2+^ ion, and Ca^2+^ release from the endoplasmic reticulum (ER) through inositol-trisphosphate receptors (IP_3_R) [15] and Raynodine receptors (RyR) [32, 14, 21], (ii) Ca^2+^ efflux through sarco/endoplasmic reticulum Ca^2+^-ATPase (SERCA) pump [31] that transfers Ca^2+^ from the cytosol of the cell to the lumen of the ER and plasma membrane Ca^2+^-ATPases (PMCA) pump [7] that transfers Ca^2+^ from the cytosol to the extracellular medium.

The impact of manipulating some of these fluxes has been studied. For example, it was previously shown that applying ryanodine, a blocker of RyR, significantly reduces the frequency of SCaLTs [32, 3], while blocking the SERCA pump to prevent reuptake into the ER by thapsigargin also diminishes SCaLTs. Additionally, applying KB-R7943 (a NCX reverse mode blocker) decreases SCaLTs frequency, highlighting its significance in generating them [3]. While these results elucidate the role of some of the fluxes in affecting SCaLTs, the contribution of other fluxes remain not fully understood, especially in the context of of how they collectively produce the fast and slow components of SCaLTs.

In this study, we adopted a computational approach by developing a stochastic spatiotemporal model (SSM) that simulates Ca^2+^ dynamics in a one-dimensional homogeneous OPC process in order to systematically dissect the contributions of different fluxes to SCaLT formation. The model provides important insights into how irregular spiking and slow baseline oscillations are generated.

## 3 Results

### 3.1 Spatiotemporal model displays slow and fast oscillations

In order to replicate the SCaLTs in OPCs [32], we developed a homogeneous stochastic spatiotemporal model (SSM) (Fig. 1A, *Materials and Methods*) of several microns of an OPC process that incorporated seven different fluxes, including those associated with SOCE, NCX, PMCA, IP_3_R, RyR, leak and SERCA. Only cytosolic Ca^2+^ concentration ([Ca^2+^]_i_) was assumed to diffuse freely. Stochasticity in the model was obtained by adding an exponentially correlated noise term, generated using an Ornstein-Uhlenbeck process, to the slow inactivation variable of IP_3_R flux. The noise process with its slow time constant was interpreted to represent the slow clustering of IP_3_R.

**Figure 1:**
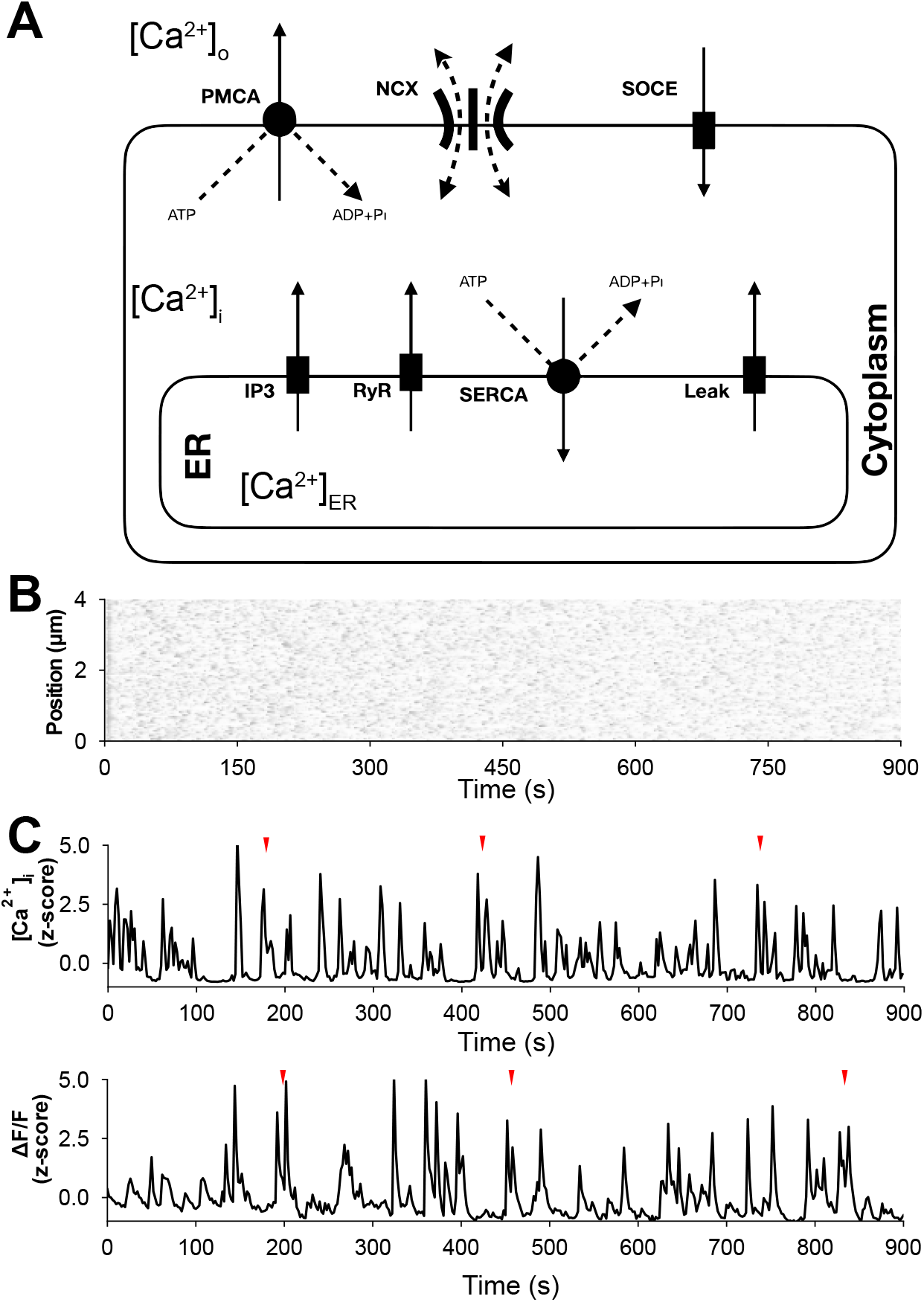
Framework and outcome of the stochastic spatiotemporal model (SSM) of spontaneous Ca^2+^ local transients (SCaLTs). (A) Schematic of the two-compartment SSM showing all fluxes across the membrane of the cell and ER included in the model. The two compartments represent cytosolic and ER Ca^2+^ concentration. (B) Spatiotemporal heatmap of Ca^2+^ amplitude over time produced by the SSM. (C) Time courses of z-scored Ca^2+^ signals at a “region of interest” along an OPC process obtained using the SSM (top) or a recorded Ca^2+^ fluorescence signal (bottom).

Simulating the SSM, using the parameter values reported in Table S1 (unless otherwise noted), over time for 900 s produced irregular spiking events distributed uniformly within the OPC process (Fig. 1B). These spikes did not appear to exhibit any specific pattern. To further examine these events, a random spatial location (“region of interest” [32]) along the process was selected and the time course of the Ca^2+^ concentration ([Ca^2+^]_i_) was then tracked (Fig. 1C, top). To allow for direct comparison with Ca^2+^ fluorescence recordings taken from rat pup OPCs and measured in terms of the ratio ΔF/F (Fig. 1C, bottom), experimental and simulated traces were both z-scored and plotted with the same sampling rate of 0.5 Hz. The resulting simulated signal was strikingly similar to the Ca^2+^ recording. They both exhibited irregular spikes superimposed on a baseline that displayed slow oscillations. The spikes had faster upstrokes than downstrokes, a hallmark of Ca^2+^ signals in OPCs, and sporadically appeared as doublets (Fig. 1C, red arrow heads). These results suggest that the combination of fluxes included in the model are sufficient for producing the intrinsic Ca^2+^ signals seen in OPCs and that the slow time scale of the stochastic IP_3_R clustering could be underlying the slow baseline oscillations.

### 3.2 Model validation and statistical analysis of SCaLTs

In order to validate the SSM against experimental recordings, simulated and experimental datasets consisting of 110 independent simulations of 900 s with a sampling rate of 0.5 Hz were used. After selecting regions of interests, the magnitude of the z-scored experimental and simulated Ca^2+^ signals were binned and histograms representing the number of points in each bin were plotted (Fig. 2A,B left). Although the profiles of experimentally obtained histograms were fairly heterogeneous, the majority of them (65%) were typically clustered around zero, with a relatively thin right-tail skew (Fig. 2B left). The parameters reported in Table S1 replicated these data very closely (Fig. 2A, left). The remaining 35% of histograms obtained from experimental recordings exhibited profiles that were roughly bimodal and centered around zero (Fig. 2B right). We surmised that varying the maximum rate of Ca^2+^ efflux from the cytosolic compartment (including *v*_PMCA_, *v*_3_) could explain the heterogeneity in the data. To recapitulate this, we assigned 35% of simulations a randomly chosen Ca^2+^ outflow rate between 50% and 100% of their default values (Fig. 2A, right). Simulations with lower efflux generated nearly bimodal distributions centered at zero, in agreement with the smaller population of experimental recordings.

**Figure 2:**
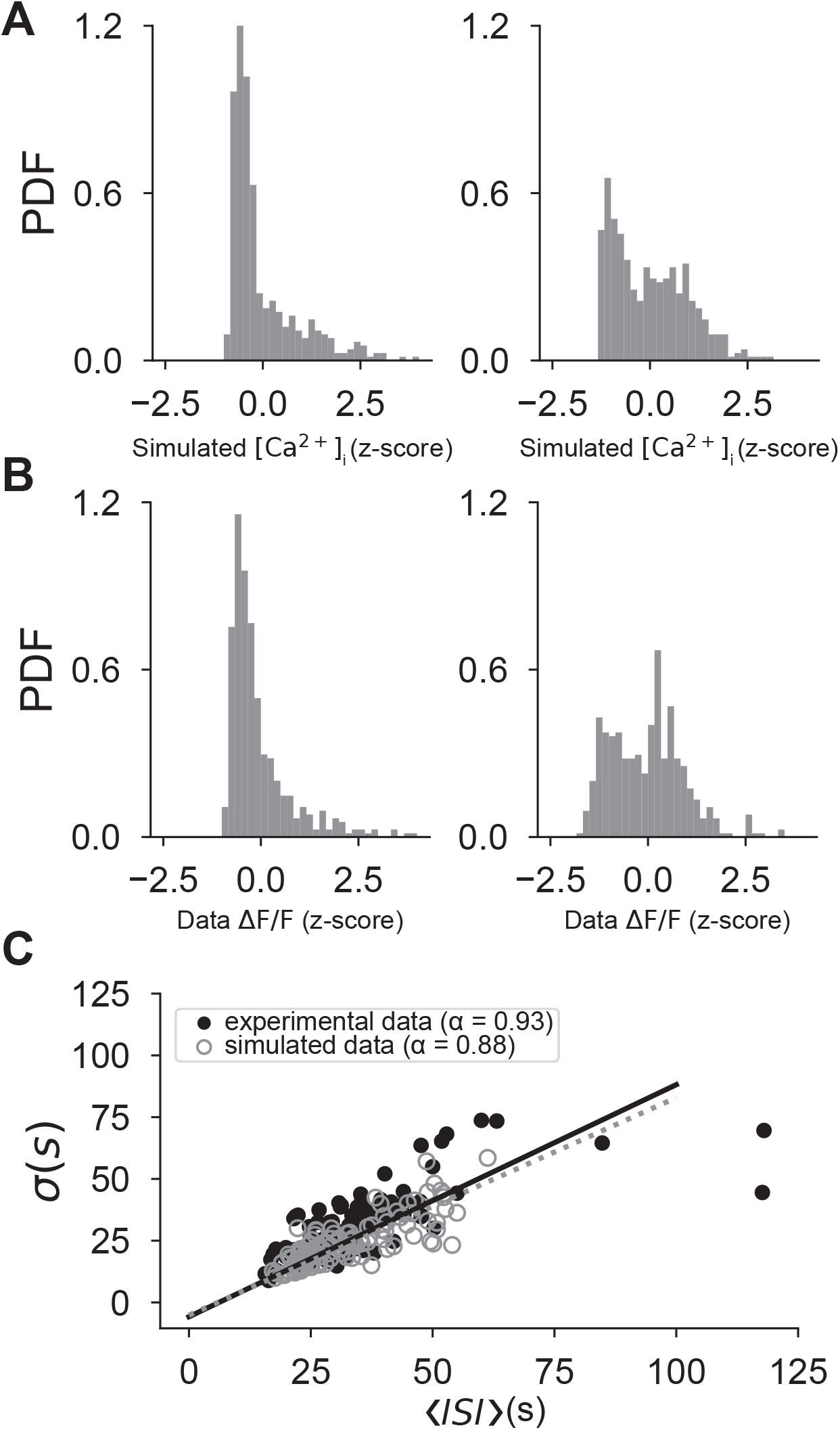
Comparing the outcomes of the SSM to experimentally recorded heterogeneous fluorescence Ca^2+^ signals. (A) Histograms of two z-scored simulated Ca^2+^ signals produced by the SSM obtained using the default parameter values listed in Table S1 (left), or after randomly reducing the maximum outward Ca^2+^ fluxes from the cytosolic compartment by 50-100% of their default values (right). (B) Histograms of two z-scored recorded Ca^2+^ signals, showcasing two typical distributions obtained from a heterogeneous data: a left-skewed one clustered around zero and seen in 65% of the recordings (left), and a bimodal one centered around zero and seen in 35% of the recordings (right). (C) Linear fits of the standard deviation of ISIs *σ* of both experimental (solid, black) and simulated (dashed, gray) recordings versus average ISIs 〈*ISI*〉, as defined by Eq. [29]. Each circle represents one single experimental (black solid) and simulated (gray open) recording.

To further explore how simulations compare to fluorescence recordings, we analyzed two key properties associated with Ca^2+^ spikes in the latter group, namely that (i) the ISI standard deviation *σ* is of the same magnitude as the average ISI *〈ISI*〉, and that (ii) the relationship between *σ* and 〈*ISI*〉 is approximately linear given by Eqs. [29] [10]. These two properties indicate that Ca^2+^ spikes are mainly stochastic events, not generated by a regular oscillatory mechanism perturbed by noise, and that their linear fit has a slope close to 1 [10]. To verify this, we plotted ISI standard deviation *σ* against average ISI 〈*ISI*〉 for both experimental and simulated recordings (Fig. 2C). The results obtained showed that these spikes were indeed stochastic in nature, that the simulations exhibited variability very similar to that seen in the data, and that the slopes of the linear fit for both datasets were very close to each other and almost equal to 1 (*α*_data_ = 0.93, *α*_sim_ = 0.88). Additionally, the intercepts for both simulation and data were almost zero, indicating there is no absolute refractory period [10]. These results thus validate the model and highlight the conclusion that information coding is actually embedded in the frequency of Ca^2+^ spikes rather than in their amplitude patterns.

### 3.3 Contributions of fluxes to SCaLTs

After model validation, we went on to explore how fluxes included in the SSM contribute to producing fast spiking and slow oscillations in [Ca^2+^]_i_ (Fig. 3A). To do this, we first tested the effects of blocking the SERCA pump by setting *J*_SERCA_ term in the SSM to zero (Fig. 3B). Our results showed that, at first, [Ca^2+^]_i_ rapidly increased as the ER depleted, then steadily decayed to zero in a manner similar to that previously seen when blocking SERCA pump with thapsigargin [3]; fast spiking and slow oscillations were both abolished in this case. We then explored the role of RyR in generating SCaLTs by setting the *J*_RyR_ term to zero in the SSM. The removal of this flux significantly reduced SCaLTs and lowered the overall baseline Ca^2+^ level (Fig. 3C); the reduced number of SCaLTs was further confirmed when computing the average frequency of these SCaLTs over multiple realizations of the model (110 in total) in the presence and absence of this flux term (inset). Such an outcome was in agreement with experimental observations obtained when blocking RyR with Ryanodine [32]. Similar results were predicted by the SSM when removing the IP_3_R flux (*J*_IP3_) and flux due to PMCA pumps (*J*_PMCA_). In the former case, the Ca^2+^ spikes and slow oscillations completely disappeared (Fig. 3D), an expected outcome in view of the fact that the stochastic term is embedded in the inactivation variable of this flux term, while in the latter case, SCaLTs were significantly reduced (Fig. 3E). Computing the average frequency of these SCaLTs over multiple realizations (110 in total) in the presence of absence of this latter flux confirmed the result (inset).

**Figure 3:**
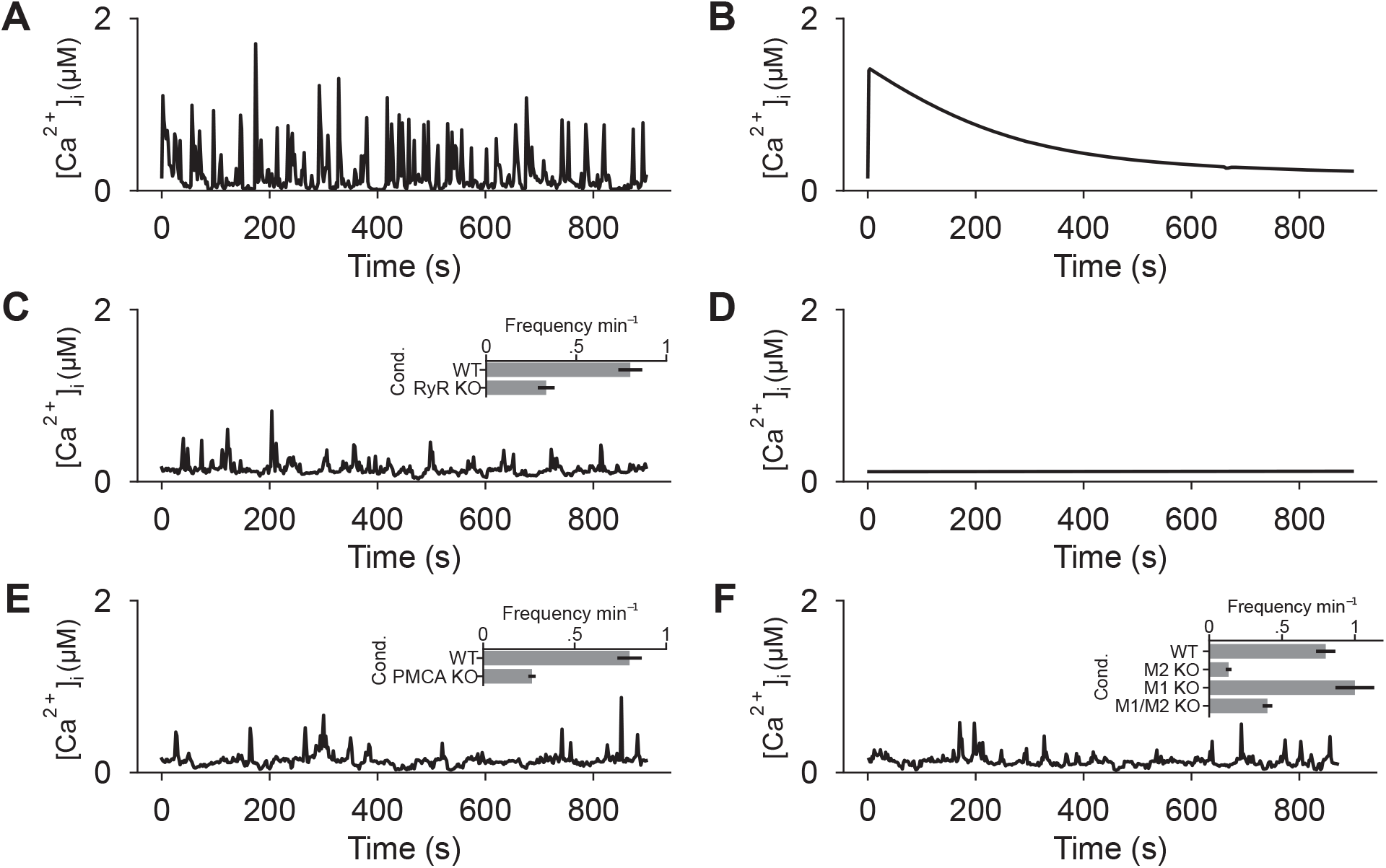
Effects of blocking Ca^2+^ fluxes on SCaLTs in the SSM. Cytosolic Ca^2+^ concentration [Ca^2+^]_i_ when (A) all fluxes were left unaltered (i.e., maximum flux rates were left at their default values – labeled wildtype), (B) maximum SERCA flux rate (*v*_3_) was set to zero, (C) maximum RyR flux rate *(v_r_*) was set to zero, (D) maximum IP_3_R flux rate (*v*_IP3_) was set to zero, (E) maximum PMCA flux rate (*v*_PMCA_) was set to zero, and (F) direct and reverse modes of NCX were blocked individually or collectively. Insets in C, E and F show average frequencies of SCaLTs obtained from 110 different simulations (realizations) of the SSM for each condition. WT = wildtype, RYR KO = RYR flux removed, PMCA KO= PMCA flux removed, M1 = mode 1 of NCX flux removed, M2 = mode 2 of NCX flux removed, None = NCX flux removed.

The NCX channel normally operates in its “forward” mode (M1), using the electrochemical gradient of Na^+^ to remove Ca^2+^ from the cell, but switches to its “reverse” mode (M2) when internal Na^+^ levels are elevated, fluxing Ca^2+^ inward. In order to study the effects of NCX flux, we removed either M1, M2 or both modes by discarding one or both terms in the numerator of Eqn. [22] (Fig. 3F). Removal of the reverse model M2 showed a significant decrease in SCaLTs, in agreement with experimental observations obtained when blocking this mode pharmacologically with KB-R7943 [3]. In contrast, the SSM predicted that the removal of the direct mode M1 increases SCaLTs due to trapping of Ca^2+^ within the cytosol and the subsequent activation of RyR and IP_3_R. Interestingly, the model also predicted that the net effect of removing both modes decreases SCaLTs but not to the same extent as when removing M2 only. These differences were further corroborated by computing the average frequency of these SCaLTs over multiple realizations (110 in total) in the presence of absence of these modes (inset).

### 3.4 Characterizing the regimes of behaviour exhibiting specific patterns of SCaLTs

To analyze the underlying dynamic of the SSM, we first simplified the stochastic spatiotemporal model into a deterministic temporal model (DTM). This was done by removing the stochastic term from the inactivation variable of IP_3_ flux (Eqn. [6]), as well as discarding the diffusion term in the cytosolic Ca^2+^ (Eqn. [1]). Using the continuation method in AUTO, we computed the bifurcation diagram of [Ca^2+^]_i_ in this simplified model with respect to *V_s_*, the maximum SOCE flux (Fig. 4A). The resulting dynamic structure was highly complex with a series of bifurcations that altered the behaviour of the model significantly.

**Figure 4:**
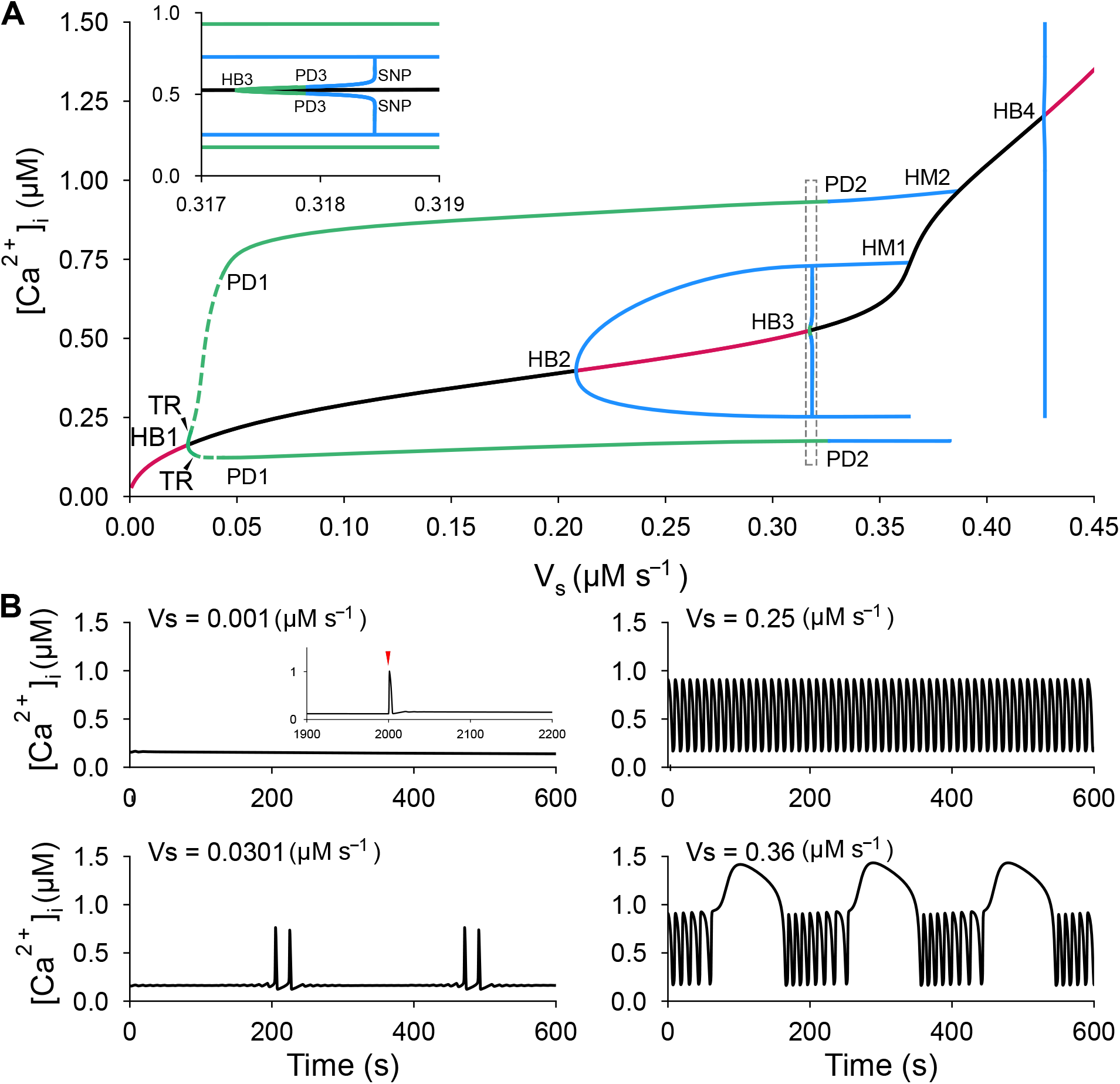
Deterministic temporal model (DTM) displays diverse Ca^2+^ signals when altering the maximum rate of SOCE flux. (A) One-parameter bifurcation of intracellular Ca^2+^ concentration [Ca^2+^]_i_ with respect to maximal SOCE conductance *V_s_*, highlighting the various regimes of behaviour. Line colours indicate branch type: magenta (black) = branch of stable (unstable) equilibria, solid/dotted green (solid blue) = envelopes of stable (unstable) periodic orbits. Bifurcations are indicated by their labels: HB = Hopf, PD = period doubling, HM = homoclinic, TR = torus, SNP = saddle-node of periodics. (B) Time series of DTM with the maximal flux rate of SOCE (*V_s_*) set to various values as reported in the legend of each panel. The following profiles have been showcased (clockwise from top/left): quiescent/excitable, tonic spiking, mixed-mode oscillations (doublets), and burst-like. Inset in the top/left panel: single spike generated by injecting a 1 s long 0.03 *μ*M Ca^2+^ pulse at 2000 s (red arrow).

Specifically, we found that for small *V_s_* (Fig. 4A), there was a branch of stable equilibria (magenta line) that lost stability at a Hopf bifurcation HB1 (black line). This Branch then regained stability at another Hopf bifurcation HB2 (magenta line) before losing it again at the Hopf bifurcation HB3 (black line) until finally regaining it at the Hopf bifurcation HB4 (magenta line). At HB1, two envelopes of stable limit cycles (solid/dotted green lines) emerged. These envelopes displayed initially torus bifurcation (TR) very close to HB1 (Fig. 4A), and then period doubling bifurcation (PD1) close to *V_s_* ≈ 0.05 *μ*Ms*s*^-1^; this was followed by another period doubling bifurcation (PD2) where the limit cycles became unstable (blue lines) before terminating at a homoclinic bifurcation (HM2). At HB2 and HB4, only unstable envelopes of periodic orbits (blue lines) emerged and terminated at homoclinic bifurcations (only HM1 is shown), whereas at HB3 (see inset in Fig. 4A), very small envelopes of stable limit cycles (green lines) emerged initially before becoming unstable (blue lines) at a period doubling bifurcations and then at a saddle-node of periodics bifurcation. According to this configuration, two key observations can be made: (i) a regime of bistability between the stable equilibria and stable envelopes of limit cycles exists between HB2 and HB3, and (ii) between HB3 and HB4, there are isolas of stable limit cycles (not shown).

To understand the implications of how these bifurcations affect dynamics, the DTM was simulated in each regime with the purpose of determining how the profiles of their time series appear. In the regime before HB1 (e.g., at *V_s_* = 0.001 *μ*Ms^-1^), the model was quiescent but excitable (Fig. 4B, top/left). Excitability was verified by showing that DTM was able to generate a single spike upon stimulation with a very brief (1 s) and small (0.03 *μM*) suprathreshold Ca^2+^ pulse (inset). This feature allowed the model to be sensitive to the Ornstein-Uhlenbeck noise process added to the slow inactivation variable of IP_3_R and caused it to produce random spiking events. In the regime between TR and PD1 (e.g., at *V_s_* = 0.0301 *μ*Ms^-1^), on the other hand, the model produced mixed-mode oscillations with two pronounced peaks in each cycle in the form of doublets (Fig. 4B, bottom/left). The fact that the default value of maximum SOCE flux (*V_s_*) is very close to where HB1 provides a rationale as to why the SSM was intrinsically able to exhibit random Ca^2+^ spikes and occasional doublets in simulated Ca^2+^ signals in the presence of noise (Fig. **??**C, top), as well as suggests that the doublets seen in experimentally recorded Ca^2+^ signals (Fig. **??**C, bottom) are an inherent aspect of Ca^2+^ spikes in OPCs. Note that all fluxes included in the DTM contributed to generating the single spikes and doublets, albeit to different levels (Fig. S1, S2).

Interestingly, two additional outcomes were also observed in other regimes of behaviour, including tonic spiking and burst like activity. Tonic spiking was seen in simulated Ca^2+^ signals when the value of maximum SOCE flux was set to *V_s_* = 0.25 *μ*Ms^-1^, a value that lies between PD1 and PD2 (Fig. 4B, top/right). Note that this behaviour could be also generated in the bistable regime provided that the DTM is initiated away from the stable equilibria. In the other regime between PD2 and HB4 (e.g., at *V_s_* = 0.36 *μ*Ms^-1^), however, Ca^2+^ signals exhibited a burst-like profile (Fig. 4B, bottom/right). Although Ca^2+^ fluxes each contributed to spiking in this profile and the tonic spiking one (Fig. S3, S4), it is important to note that, experimentally, these two profiles were likely non-physiological and would require a major level of SOCE potentiation to produce them.

### 3.5 Excitability and mixed-mode oscillations underpin spiking in Ca^2+^ signals produced by OPCs

In the previous section, we outlined the two most physiologically-relevant dynamic behaviours associated with SCaLTs in OPCs, namely, single spiking due to excitability and mixed-mode oscillations with doublets (Fig. 4). Here, we explored how these regimes define dynamics in the SSM, especially the source of irregularity seen in Ca^2+^ spiking.

To do this, we fist extended our previous analysis of the parameter regimes into performing a two-parameter bifurcation with respect to the maximum flux rates of IP_3_R (*ν*_IP3_) and SOCE (*V_s_*) (Fig. 5A). This was done by tracing all the Hopf bifurcations (obtained in Fig. 4A) to generate the boundaries of the various regimes of behaviour in this two-parameter space. The goal was to investigate the collective contribution of these two fluxes in defining DTM dynamics and to explore how perturbing them could alter dynamics of the SSM.

**Figure 5:**
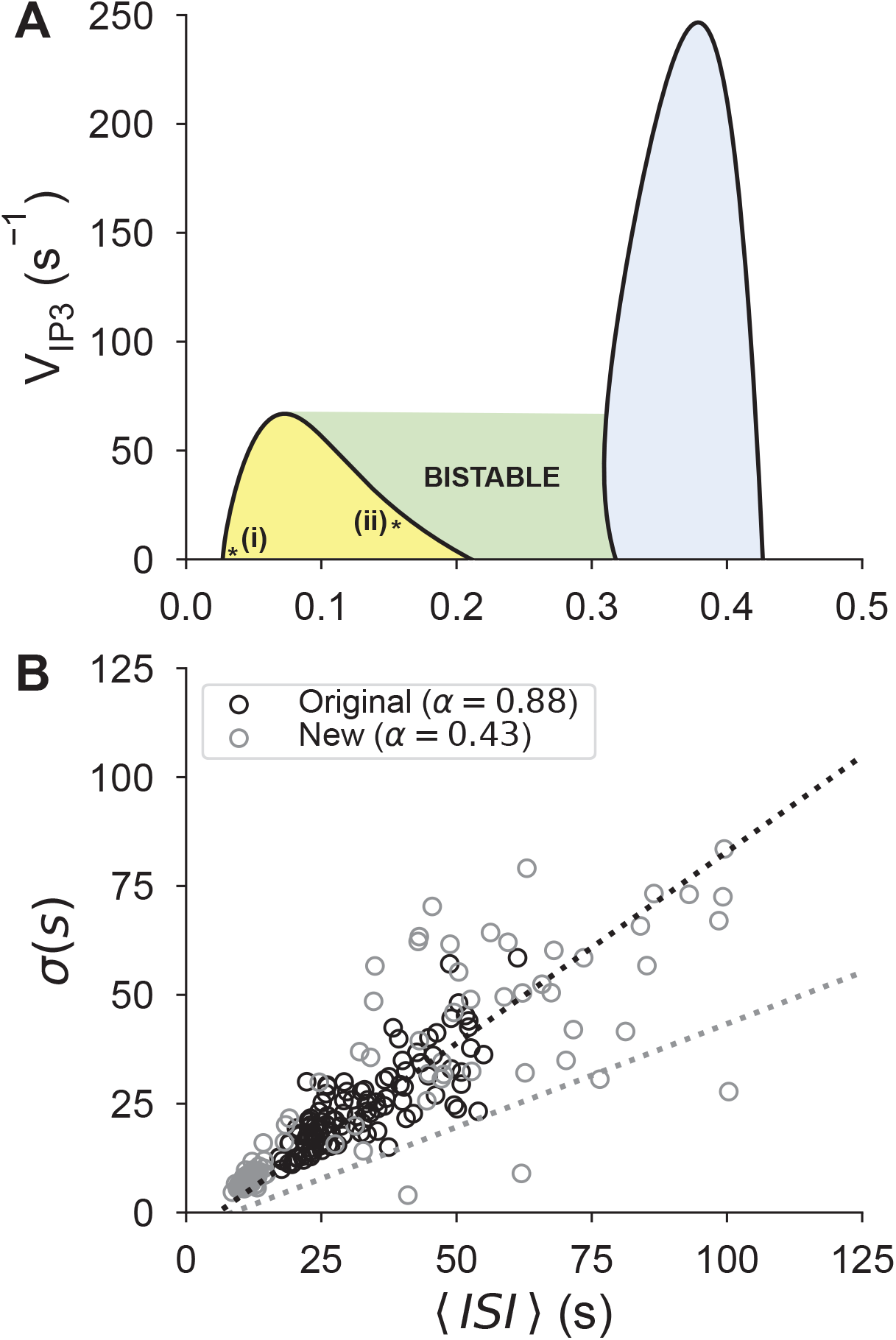
Parameter regimes defined by the DTM dictate the outcomes of the SSM, including irregular Ca^2+^ spiking and doublets. (A) Two-parameter bifurcation with respect to the maximum flux rates of IP_3_R (*v*_IP3_) and SOCE (*V_s_*) obtained by tracing the Hopf bifurcations of Fig. 4 while changing these two parameters. Four different regimes of behaviour were identified: quiescent (white), oscillatory containing mixed-mode oscillations and tonic spiking (yellow), bistable between quiescent and tonic spiking (green) and oscillatory containing tonic spiking and burst-like activity (blue). (B) Linear fits of the standard deviation of ISI’s *σ* versus average ISIs 〈*ISI*〉, as defined by Eq. [29], for two simulated datasets: an original one generated using the default parameter values of *ν*_IP3_ and *V_s_* (black), identified by the i-labeled dot in A (see Table S1), and a new one generated by setting the values of these two parameters to *ν*_IP3_ = 20 s^-1^ and *V_s_* = 0.155 *μ*Ms^-1^ (gray), identified by the ii-labeled dot in A. Dotted lines represent the linear fits, while open circles represent simulated recordings.

The resulting two-parameter bifurcation diagram obtained (Fig. 5A) showed that there were four distinct regimes, two of which were oscillatory (yellow and blue), one was quiescent (white) and one that included both, i.e., was bistable (green). The yellow-colored oscillatory regime included two types of oscillatory behaviours (Fig. 4B): mixed-mode oscillations with doublets close to the left boundary (i.e., in the vicinity of the i-labeled dot) and tonic spiking close to the right boundary (i.e., in the vicinity of the ii-labeled dot). The blue-colored oscillatory regime, on the other hand, included the tonic spiking and burst-like activity (Fig. 4B). Finally, the bistable regime included the quiescent and tonic firing activity. Similar outcomes were also obtained when plotting the two-parameter bifurcation with respect to maximum RyR *(V_r_*) and SOCE (*V_s_*) flux rates (Fig. S5)

Although it was difficult to pinpoint the boundary of the excitable regime, the white-colored region to the left of the yellow-colored oscillatory regime happened to be excitable (Fig. 5A). The default values of the IP_3_R and SOCE maximum flux rates lie very close to boundary of this region (at the i-labeled dot in Fig. 5A). These default values allowed the SSM to generate a slope close to 1 for the linear fit of *σ* versus 〈*ISI*〉 (Fig. 2C), as per Eqn. [29]. We postulated that moderately perturbing these two parameters by choosing values close to the right boundary of the yellow-colored oscillatory regime (at the ii-labeled dot in Fig. 5A) would significantly alter the slope of the linear fit. To test this hypothesis, we generated 110 independent simulations of 900 s with a sampling rate of 0.5 Hz using this new parameter combination for IP_3_R and SOCE maximum flux rates and performed the same linear fit defined by Eqn. [29] (Fig. 5B). The slope of this linear fit (gray dotted line) obtained from these new model realizations (gray open circles) was 0.43, a value substantially lower than the 0.88 slope of the linear fit (black dotted line) obtained from the original model realizations (black open circles) when default parameter values were used (the same dataset as in Fig. 2C). These results suggest that the combination of excitability and stochasticity are the necessary ingredients to generate irregularity in Ca^2+^ spikes and that the mixed-mode oscillations are causing the system not only to exhibit doublets but also to make the slope of the linear fit slightly smaller than 1 due to its periodicity.

## 4 Discussion

Spontaneous Ca^2+^ local transients (SCaLTs) in OPCs result from a complex interaction of multiple Ca^2+^ fluxes that are produced or modulated by several ion channels, receptors and pumps. These SCaLTs, through their coupling with the golli protein pathway in OPCs, suggest that they might play a key role in the development of myelin [29, 28]. In this study, we focused on the following key fluxes generated by those on the membrane of the cell: store-operated Ca^2+^ entry (SOCE), Na^+^ /Ca^2+^ exchangers (NCX) and plasma-membrane Ca^2+^ ATPase (PMCA) pumps, as well as those on the membrane of the ER: inositol-trisphosphate receptors (IP_3_R), Ryanodine receptors (RyR), sarco/endoplasmic reticulum Ca^2+^ ATPase (SERCA) pumps and leak. In this study, we used data-driven computational modeling approach to investigate how these fluxes interact together to generate the SCaLTs seen OPCs.

The approach comprised of developing a stochastic spatiotemporal model (SSM) that included all the relevant fluxes and validating the model against Ca^2+^ fluorescence data measured in terms of the ratio AF/F_0_ [32]. The SSM was implemented over a one-dimensional OPC process and incorporated an Ornstein-Uhlenbeck noise process that possessed a slow time constant into the inactivation variable of IP_3_R flux. The latter was included due to the stochastic nature of the Ca^2+^ signal and it was interpreted to represent the slow clustering of IP_3_R. The model was then used to determine the contribution of each flux to SCaLTs and to decipher their underlying dynamics, making important predictions about how the irregular spikes and slow baseline oscillations of the signal were generated.

After validating the model against data, we concluded from the study that all fluxes could reduce the frequency of SCaLTs to different levels, except for the forward mode of NCX which did the opposite. The model suggests that the trapping of Ca^2+^ inside the cell when blocking the forward mode of NCX was responsible. We showed, using the deterministic temporal model (DTM), that the random nature of SCaLTs was a result of the excitable nature of the system in the most physiological regime and that doublets were intrinsic to the system produced by noise-induced shifts to relevant parameter regimes. Using bifurcation analysis, we determined how the system depended on its current state in allowing a given Ca^2+^ flux to exert its influence and demonstrated that the irregularity in spiking would not persist if dynamics were shifted deep into the oscillatory regimes.

By plotting the linear fit of the standard deviation of interspike intervals (ISIs) of both experimental and simulated recordings against the average ISIs using Eq. [29], we demonstrated that the slope of the line was close to 1 and the intercept close to 0, indicating a highly stochastic process with no absolute refractory period. This also revealed that at its default parameter values (Table S1), the model could produce irregular Poissonian spikes and that information coding was embedded in spike frequency rather than spike amplitude [36, 10]. However, the slope of the linear fit became significantly smaller than 1 when the model was simulated deeper in the tonic spiking regime. This indicate that there is a lower level of randomness in spiking, and that excitability is the main driver of random spiking in OPCs.

Due to the stochastic nature of SCaLTs, we could not fit the model directly to the data. Indeed, experimental recordings display a great deal of statistical heterogeneity, thereby making it difficult to form any single metric to compare data and simulations. We resolved this issue by employing two approaches: one that compared the distributions of the experimental signals to those obtained by simulating the SSM (focusing on two representative distributions), and another that compared the linear fits. The parameter set in Table S1 effectively replicate the main characteristic features of the majority of the data using the two approaches. By doing so, we show that scaling the total efflux from cytosol is necessary to replicate the the bimodal distribution seen in 35% of the experimental recordings.

From a mathematical perspective, the dynamic structure of the DTM was not fully explored. There are likely interesting dynamics occurring at the regime bounded by the period doubling bifurcation point PD2 and the Hopf bifurcation HB4 (Fig. 4A). We presumed that this regime was likely occupied by infinite number of isolas, each possessing a stable envelope of periodic orbits with specific number of spikes associate with the burst-like activity. One can further investigate this behaviour and apply slow-fast analysis [11] to determine the mechanism underlying this bursting behaviour.

The spatiotemporal model in this paper consists of one homogeneous process extending in space. The model could be further improved by the addition of coupled branches to mirror the dendritic nature of OPC processes. Moreover, the ER was assumed to be uniform across the process and in close proximity to the plasma membrane. One can develop a more realistic model by including a diffusion term into Eqn. 2. Although this may provide an accurate framework to describe Ca^2+^ dynamics in OPCs, we do not expect this additional complexity to alter to conclusions of this study.

Given that myelination in the central nervous system is likely coupled to the neuronal activity of nearby cells, developing this flux-balance based stochastic spatiotemporal model in this study can provide an interface that can be further extended and analyzed to gain new insights about the intrinsic Ca^2+^ transients underpinning neuro-oligo signaling. The investigation of this topic is critical for understanding myelin plasticity and the development of treatments for leukodystrophies and demyelinating diseases.

## 5 Methods

### Experimental data

Ca^2+^ fluorescence data was obtained from rat OPCs (for more details about how such data were collected, see [32]). Briefly, cerebral cortices from postnatal day 0-2 rat pups were dissected and mechanically dissociated, with the final culture containing approximately 85%-90% OPCs. The cultured OPCs were induced to differentiate and immunostained to allow Ca^2+^ to be tracked at several sites on one cell. In total 110 time series of ΔF/F_0_ Ca^2+^ fluorescence data were obtained in 4 groups of 27 independent recordings.

### Mathematical model

We developed a flux-balanced based spatiotemporal Ca^2+^ spiking model to simulate intrinsic Ca^2+^ dynamics of OPCs. The model is a combination of (1) the Li-Rinzel IP_3_R/SERCA model of Ca^2+^ handling [22], (2) the Levine-Keizer RyR model [16], (3) the Croisier et al. SOCE model [8], and (4) the Weber et al. NCX model [38]. It consists of seven equations, with one incorporating a second-order spatial term to represent cytosolic Ca^2+^ diffusion, and another incorporating noise to provide random input to the slow inactivation variable of IP3 flux.

Descriptions and values for all the parameters of the model are given in Table S1. The equations describing the dynamics of Ca^2+^ concentration in the cytosol ([Ca^2+^]_i_) and ER ([Ca^2+^]_ER_) are given by

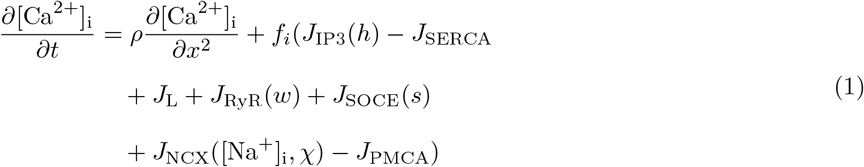

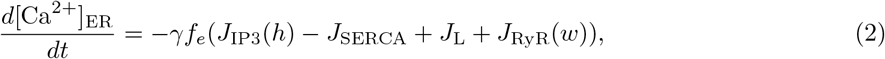

Flux due to IP_3_R (*J*_IP3_) is given by

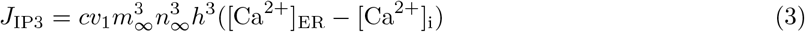

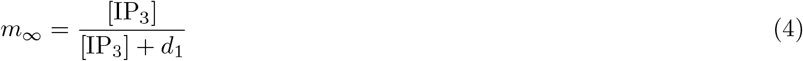

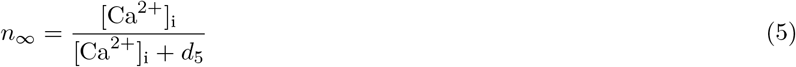

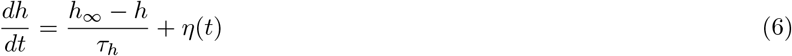

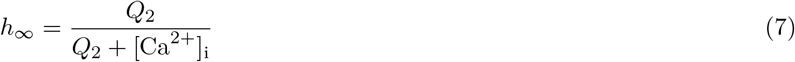

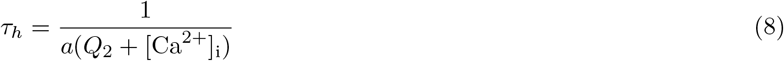

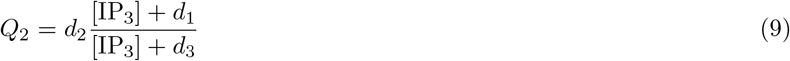

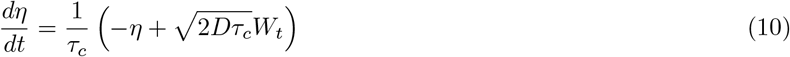

where *m*_∞_ and *n*_∞_ are the steady state activation functions (indicating fast receptor activation), and *h*(*t*) is the slow inactivation variable with an exponentially correlated noise term *η*(*t*) added it in a manner similar to that used in [20, 26]. The noise *η*(*t*) was generated by an Ornstein-Uhlenbeck process to represent slow clustering of IP_3_R necessary for reproducing the slow oscillations in Ca^2+^ signals. The parameter *τ_c_* is the characteristic correlation time of the noise, *D* is the noise intensity and *W_t_* is a Gaussian white noise process with *μ* = 0 and *σ* = 1.

Flux due to SERCA pumps ( *J*_SERCA_) is given by

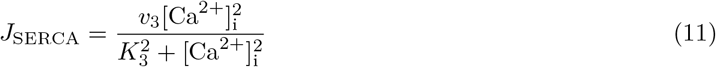

The constant Ca^2+^ leak going into the cytosol from the ER through unspecified channels (*J*_L_) is given by

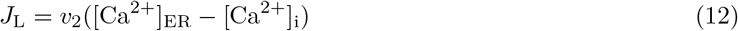

The Ca^2+^ flux due to RyR (*J*_RyR_) is given by

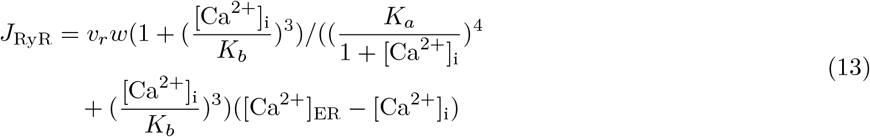

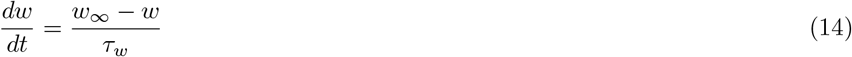

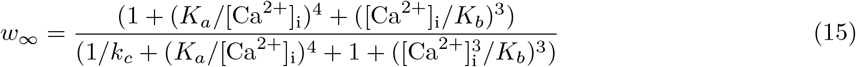

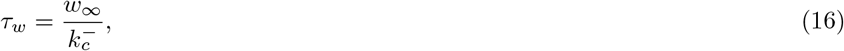

where *w*(*t*) is a dynamic slow inactivation variable.

The flux of Ca^2+^ through the SOCE channel (*J*_SOCE_) is given by

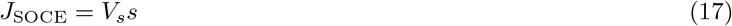

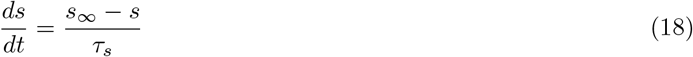

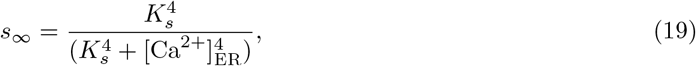

where *s*(*t*) is a slow activation variable.

The reversible, polarization-dependant, Ca^2+^ flux between the cytosol to the extracellular medium via the NCX exchanger (*J*_NCX_) is given by

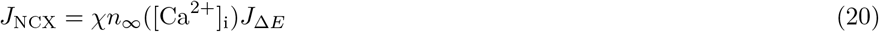

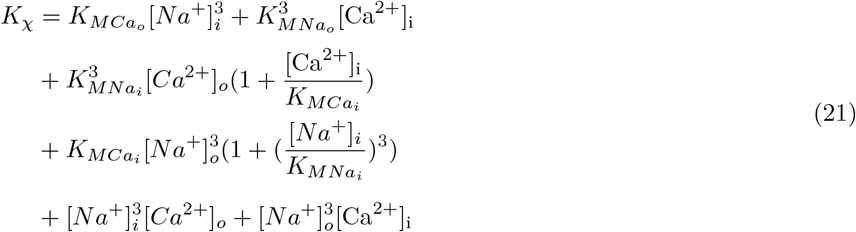

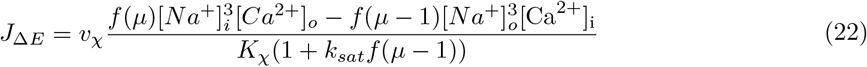

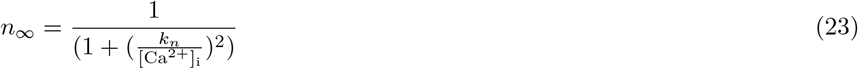

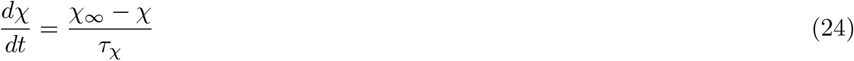

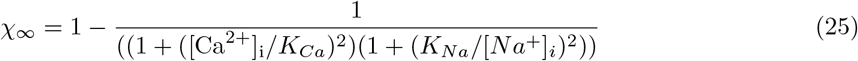

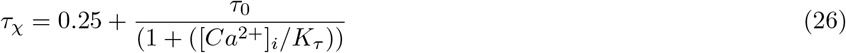

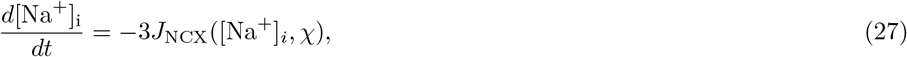

where *χ*(*t*) is the slow inactivation variable of the NCX and 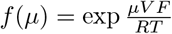.

Finally, Ca^2+^ flux due to PMCA pump (*J*_PMCA_) is given by

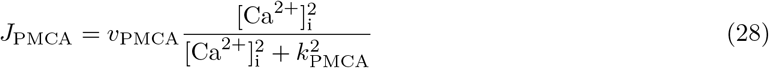

To solve this system of differential equations, we implemented the Forward Time Centered Space (FTCS) scheme with Neumann boundary conditions 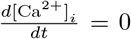 at both ends. The Ornstein-Uhlenbeck equation was solved using the Euler-Murayama scheme. The time was discretized in intervals of 0.01 s and space in intervals of 0.01 μm.

### Data analysis

Spontaneous Ca^2+^ local transients (SCaLTs) in both simulated and experimental data were identified by detecting peaks greater than 20% from the baseline, or 1 standard deviation above the mean. For all analyses, comparing simulated and experimental data time series were z-scored. To analyze the variability of Ca^2+^ signals, we applied the method of Dupont et al. [10]. For each realization of both data and simulation, the interspike interval standard deviation *σ* was plotted against the mean interspike interval *T*_avg_ and fitted with the linear equation

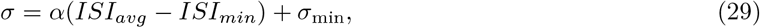

where *ISI_min_* and *σ_min_* are the minimum interspike interval and standard deviation, respectively. The intercept of the equation corresponds to an absolute refractory period in which no more spikes can occur.

### Hardware & Software

Computer simulations of the SSM and DTM Ca^2+^ models were run on a workstation with an Intel CORE i7 12700k running at 3.6 Ghz, and 32 GB of DDR5 RAM. Solutions to the SSM system were computed using a custom Python script optimised with the Numba JIT compiler [19]. Versions of the SSM solver code are available for download at https://github.com/loprea91/Ca2SSM in Python, MATLAB, and Julia. The deterministic reduced system was numerically solved in python; XPPAUT (a freeware available online at http://www.math.pitt.edu/~bard/xpp/xpp.html) was used to compute the bifurcation diagrams. The DTM code and XPPAUT function file are also available for download.

This work was supported by the Natural Sciences and Engineering Research Council of Canada (https://www.nserc-crsng.gc.ca/index_eng.asp) discovery grants (numbers: 5013485-2019, 341534-2012), and by the National Institutes of Health (https://www.nih.gov/) (number: GM083889).

