## Supplemental Table 1 and Figures S1-S5 for "Characterizing the role of Ca^2+^ fluxes in defining fast and slow components of spontaneous Ca^2+^ local transients in OPCs using computational modeling"

Table S1: Parameter values used in the SSM and DTM model simulations

| Symbol | Value | Description |
|---|---|---|
| | | $IP_3$ flux |
| $c$ | 0.111 | (ER vol)/(cytosolic vol) |
| $v_{\mathrm{IP3}}$ | 0.88 $s^{-1}$ | Max channel flux |
| $[IP_3]$ | 0.2 µM | $IP_3$ concentration |
| $d_1$ | 0.13 µM | Dissociation constant $IP_3$ |
| $d_5$ | 0.08234 µM | Dissociation constant $Ca^{2+}$ (activation) |
| $a_2$ | 0.2 s | Binding constant $Ca^{2+}$ (inhibition) |
| $d_2$ | 1.049 µM | Dissociation constant $Ca^{2+}$ (inhibition) |
| $d_3$ | 0.9434 µM | Dissociation constant $IP_3$ |
| | | SERCA flux |
| $v_3$ | 120 µM $s^{-1}$ | Maximal rate of SECRA pump |
| $K_3$ | 0.3 µM | Dissociation constant for SERCA |
| | | Leak flux |
| $v_2$ | 0.5 $s^{-1}$ | Rate constant for store leak |
| | | RyR flux |
| $v_r$ | 18 $s^{-1}$ | Rate constant for RyR |
| $K_a^4$ | 0.0192 µM $^4$ | Ratio of kinetic constants |
| $K_b^3$ | 0.2573 µM $^3$ | Ratio of kinetic constants |
| $K_c$ | 0.0571 | Ratio of kinetic constants |
| $k_c^-$ | 0.1 $s^{-1}$ | kinetic constants |
| | | SOCE flux |
| $V_s$ | 0.0301 µM $s^{-1}$ | SOCE maximum flux |
| $K_s$ | 50 µM | STIM ER $Ca^{2+}$ affinity |
| $\tau_s$ | 30 s | SOCE timescale |
| | | NCX flux |
| $[Ca^{2+}]_o$ | 12 µM | Calcium concentration outside of the cell |
| $[Na^+]_o$ | 14 mM | Sodium concentration outside of the cell |
| $v_{\mathrm{NCX}}$ | 1.4 A/F | NCX maximal flux |
| $\mu$ | 0.35 | Position of the energy barrier |
| $V$ | -80 mV | Transmembrane potential |
| $F$ | 96.5 mC/mol | Faraday's constant |
| $R$ | 8.314 $JK^{-1}mol^{-1}$ | Gas constant |
| $T$ | 300 K | Temperature |
| $k_{sat}$ | 0.25 | Saturation factor at negative potential |
| $K_{MCa_o}$ | 1.3 mM | Extracellular $Ca^{2+}$ dissociation constant |
| $K_{MNa_o}$ | 97.63 mM | Extracellular $Na^+$ dissociation constant |
| $K_{MCa_i}$ | 12.3 mM | Intracellular $Ca^{2+}$ dissociation constant |
| $K_{MNa_i}$ | 0.0026 mM | Intracellular $Na^+$ dissociation constant |
| $K_n$ | 0.1 µM | Calcium activation |
| | | PMCA flux |
| $K_{\mathrm{PMCA}}$ | 0.8 µM | Dissociation constant for PMCA pump |
| $v_{\mathrm{PMCA}}$ | 0.6 µM $s^{-1}$ | Maximal rate of PMCA pump |
| | | Ornstein-Uhlenbeck process |
| $\tau_c$ | 0.01 s | Characterisitc time constant |
| $D$ | 1.2 | Noise intensity |
| | | Other parameters |
| $f_i$ | 0.01 | Fraction of free calcium in cytosol |
| $f_e$ | 0.025 | Fraction of free calcium in ER |
| $\gamma$ | 9 | Ratio of cystolic to ER volume |
| $\rho$ | 5e$^{-5}$ µM $s^{-1}$ | Diffusion coefficient |

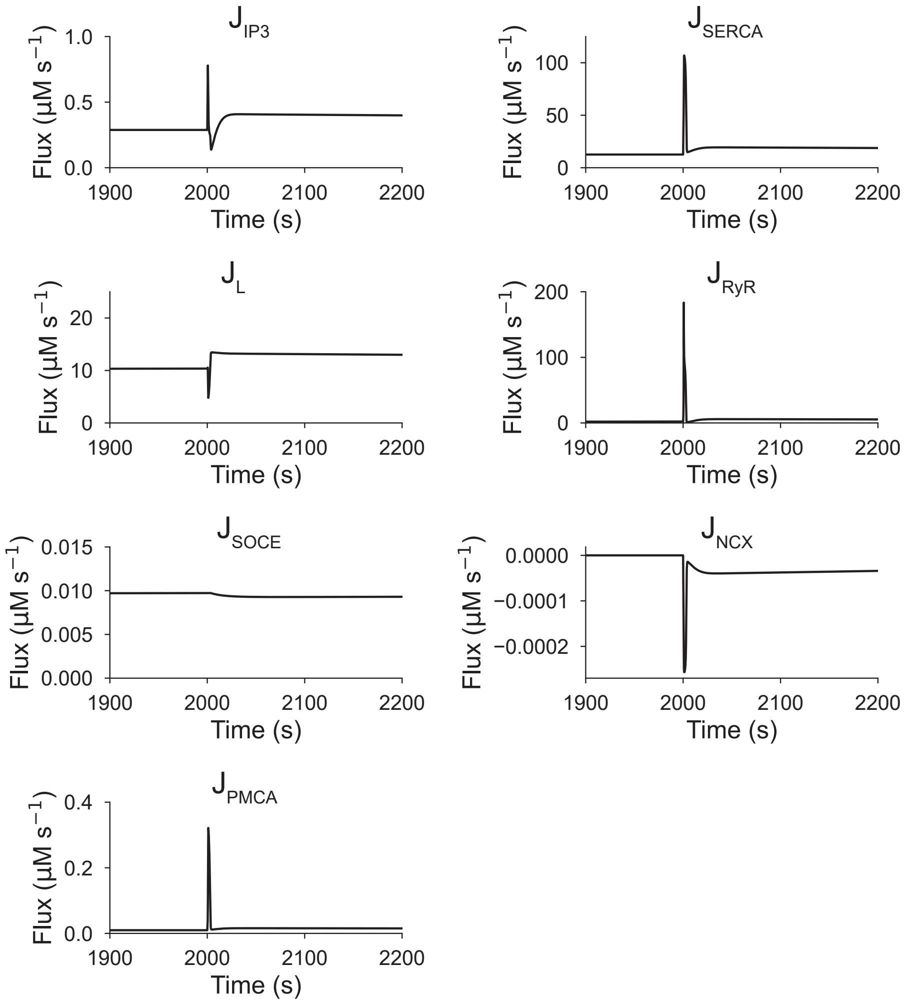


Figure S1: Simulated time courses of all of the $Ca^{2+}$ fluxes included in the deterministic temporal model (DTM) when $V_s = 0.001\ \mu\text{Ms}^{-1}$. The simulations show the contribution of each flux to generating a single spike upon stimulation with a 1 s long 0.03 $\mu$M $Ca^{2+}$ pulse at 2000 s.

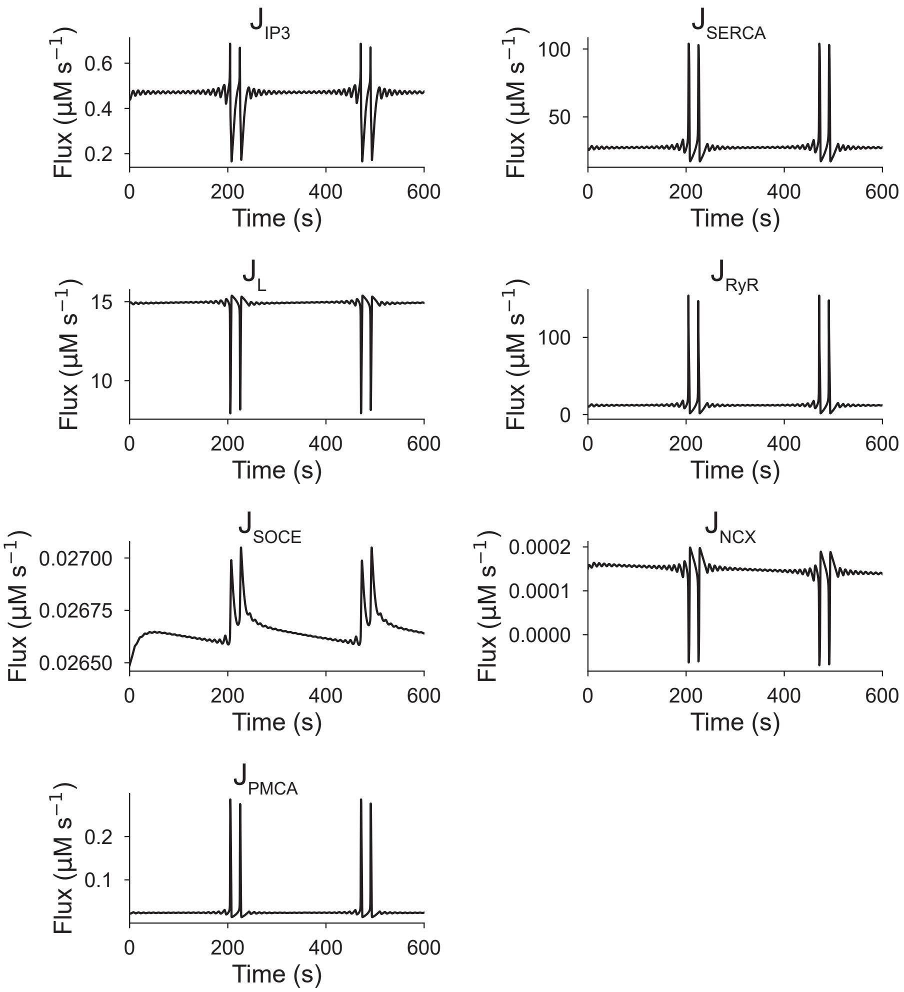


Figure S2: Simulated time courses of all of the $Ca^{2+}$ fluxes included in the deterministic temporal model (DTM) when $V_s = 0.0301$ $\mu$Ms$^{-1}$. The simulations show the contribution of each flux to the formation of doublets in the mixed-mode oscillation regime.

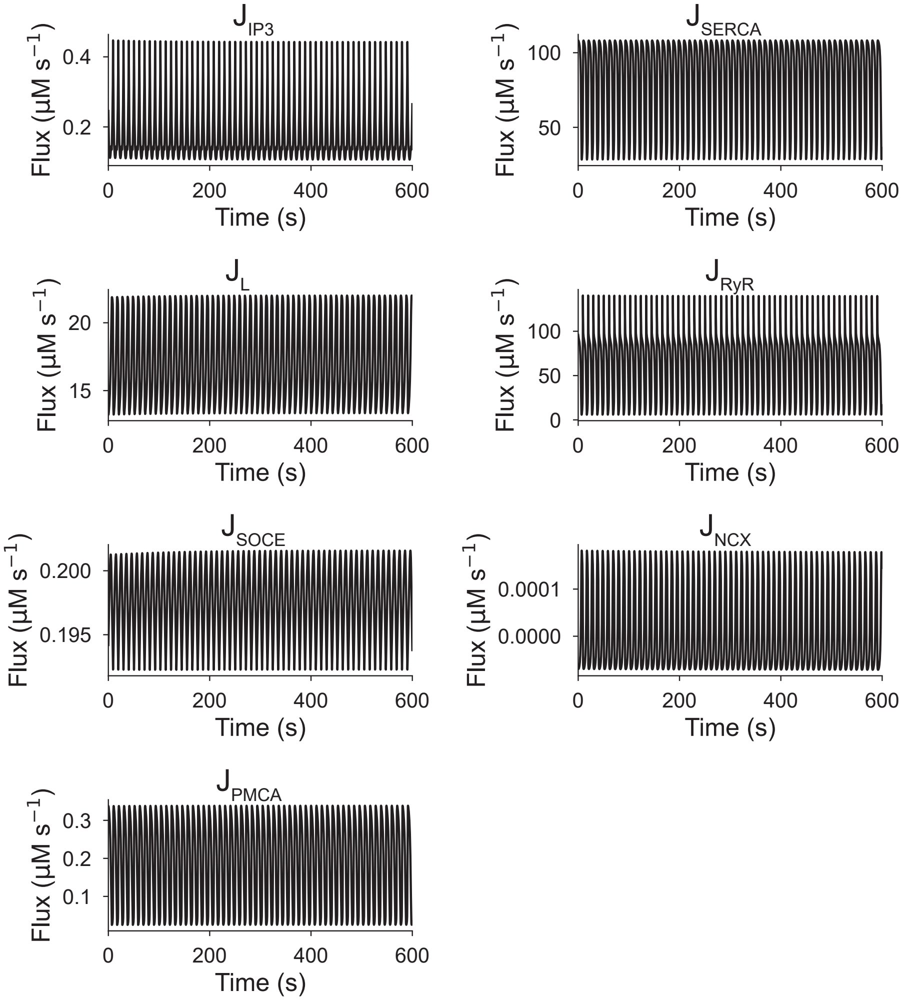


Figure S3: Simulated time courses of all of the $Ca^{2+}$ fluxes included in the deterministic temporal model (DTM) when $V_s = 0.25$ $\mu$Ms$^{-1}$. The simulations show the contribution of each flux to generating tonic spiking.

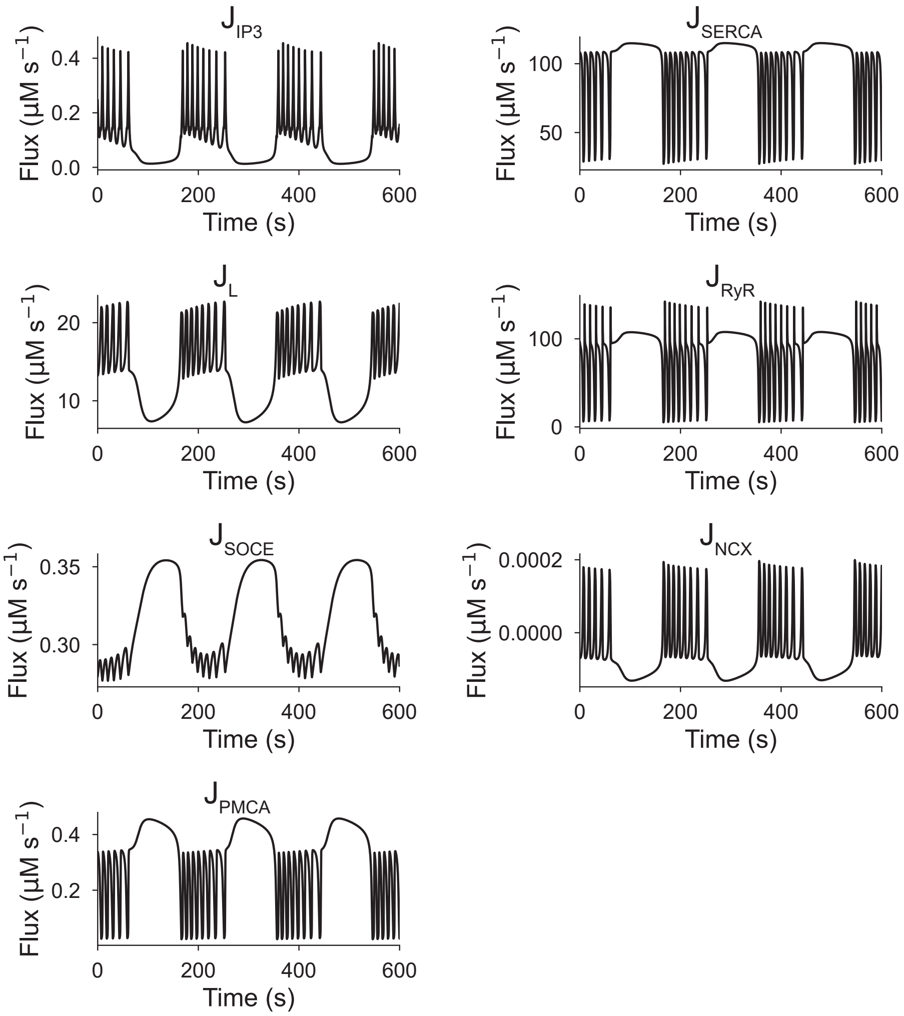


Figure S4: Simulated time courses of all of the $Ca^{2+}$ fluxes included in the deterministic temporal model (DTM) when $V_s = 0.36\ \mu\text{Ms}^{-1}$. The simulations show the contribution of each flux to generating burst-like activity.

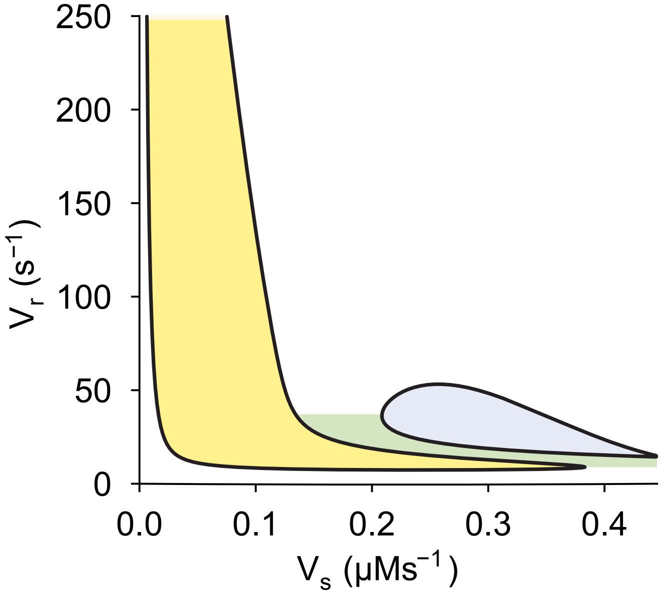


Figure S5: Two-parameter bifurcation with respect to the maximum flux rates of RyR ($V_r$) and SOCE ($V_s$) obtained by tracing the Hopf bifurcations of Fig.4A while changing these two parameters. Four different regimes of behaviour were identified: quiescent (white), oscillatory containing mixed-mode oscillations and tonic spiking (yellow), bistable between quiescent and tonic spiking (green) and oscillatory containing tonic spiking and burst-like activity (blue).
